# From prediction to engagement: defining tumor-reactive T cells through biological interaction

**DOI:** 10.64898/2026.05.31.729129

**Authors:** Nicholas P. Restifo, Suman K. Vodnala, David B. Thompson, Nicholas D. Klemen, Ira Mellman, Denitsa Milanova

## Abstract

Adoptive T-cell therapy (ACT), particularly tumor-infiltrating lymphocyte (TIL) therapy, demonstrates that durable regression of solid tumors can occur when polyclonal T cells target authentic tumor antigens, including private neoantigens. Yet scalable identification of tumor-reactive clonotypes remains limited because the relevant cells are rare, patient-specific, and poorly captured by indirect markers, expansion behavior, or predictive algorithms. We examined whether productive engagement between T cells and tumor cells or antigen-presenting cells provides a more direct organizing principle for tumor-reactive T-cell discovery. Across published tumor and blood datasets reanalyzed here, T cells recovered through physical clustering, target-cell extraction, or antigen-dependent activation are enriched for clonotypes occupying antigen-experienced, progenitor-like states with restrained cytotoxic differentiation. Productive engagement can therefore serve as a practical enrichment variable, enabling recovery of rare neoantigen-reactive T cells from circulation while preserving developmental states associated with persistence and therapeutic responsiveness. These analyses support a shift in tumor-specific T-cell discovery from prediction-first nomination to interaction-first recovery: enrich for T cells bearing evidence of prior tumor-antigen encounter in vivo, recover clonotypes preserved in a restrained progenitor-like state, expand them before culture competition erases them, and reinforce them therapeutically with matched antigenic information.

## Introduction

For more than three decades, adoptive T-cell therapy has established a central principle in cancer immunology: tumor-reactive T cells can mediate deep and sometimes durable regression of treatment-refractory solid tumors [1,2]. These responses are thought to reflect prior in vivo encounter with tumor-derived antigen, but not necessarily direct first contact with a malignant cell. Initial priming may occur through cross-presenting dendritic cells in tumor-draining lymph nodes, whereas continued antigen encounter and state imprinting can occur in the tumor through interactions with malignant cells and intratumoral antigen-presenting cells [3,4]. In such cases, the immune system has already solved the problem of tumor antigen reactivity. However, generalizing this biological fact into a scalable method for identifying, recovering, and amplifying tumor-reactive clones at therapeutically meaningful numbers remains unsolved, because these tumor-relevant antigens are unique in virtually every patient. This is the central translational bottleneck for solid-tumor ACT: not whether tumor-reactive T cells exist, but whether they can be reproducibly selected, expanded, and maintained before irrelevant or terminally differentiated cells dominate the highly personalized, patient-specific product.

Most modern cellular immunotherapies are built on an instructional model: nominate the target first, then engineer, elicit, or expand T cells against it. This strategy works when the target is shared, abundant, safe, and biologically decisive. It becomes fragile when the relevant antigens are private, sparsely presented, and embedded in patient-specific immune histories. Investigators nominate candidate targets—through surface-marker biology, tumor sequencing, or computational prediction—and then engineer or elicit T cells against those predefined antigens. CAR-T cells, TCR-engineered T (TCR-T) cells, and most cancer vaccines all follow this logic. The model has produced extraordinary successes, especially in hematologic malignancies, but those successes arose under unusually favorable biological conditions.

CD19- and BCMA-directed CAR T cells succeeded in settings where malignant cells express abundant lineage-restricted targets and where on-target injury is clinically manageable [5–11]. Epithelial carcinomas, which account for most adult malignancies, rarely offer analogous targets [12–15]. Their candidate surface antigens are often heterogeneous, shared with essential normal tissues, or merely overexpressed rather than tumor exclusive [16]. TCR-T cell approaches can reach intracellular and mutation-derived epitopes [17–20], but their usefulness remains constrained by HLA restriction, the presence of the cognate mutation-derived peptide, and the rarity of any given actionable pMHC target [21–26]. Thus, the hematologic CAR-T template has not failed to extend to solid tumors because of insufficient engineering alone; it has failed because most solid cancers do not provide the same kind of target.

To address the limitations of shared tumor-associated antigens, the field has moved toward increasingly precise, patient-specific targets [27]. Neoantigens represent the most refined version of this instructional strategy, because they are derived from somatic mutations and can, in principle, focus immunity on tumor-restricted determinants. Personalized neoantigen vaccination has demonstrated that patient-specific tumor mutations can be used to generate broad neoantigen-specific T-cell immunity [28–30]. In surgically resected pancreatic cancer, studies of autogene cevumeran extended this principle to an individualized RNA platform, showing de novo priming of high-magnitude, poly-epitopic, durable CD8+ T-cell responses [30,31].

However, these studies were performed largely in adjuvant or minimal-residual-disease settings and do not by themselves establish that vaccination can reliably eliminate bulky established tumors. In the framework proposed here, the key therapeutic unit is not the predicted neoantigen alone, nor the vaccine-induced response alone, but the matched pair: a patient-specific antigenic representation used to recover and reinforce T cells that have already shown evidence of tumor-antigen engagement.

Thus, the central constraint remains: specifying an antigen is not the same as demonstrating tumor recognition. A candidate neoantigen must be expressed, processed, presented by the relevant HLA molecule, and displayed at sufficient density on tumor cells. The induced T cells must then have sufficient functional avidity and an appropriate differentiation state to recognize antigen in that native tumor context. Classical peptide-vaccine studies illustrate this distinction. Vaccine-elicited CD8+ T cells can recognize exogenous peptide or vaccine-loaded APCs while showing limited recognition of naturally processed antigen and inefficient tumor-cell lysis. By contrast, endogenous tumor-associated T cells can exhibit higher recognition efficiency and more effective tumor lysis [32–35]. Even successful vaccination may therefore generate measurable immunity without ensuring engagement with the targeted cell itself [36]. This limitation is not an argument against vaccines, but it defines their boundary: vaccines can instruct and amplify; they do not, by themselves, verify that the induced or expanded T cells have recognized tumor in vivo.

These limitations point to a different approach: rather than specifying tumor antigens in advance, the problem can be inverted by recovering the T cells that have already solved it in vivo. Tumor-infiltrating lymphocytes provide a different form of evidence. Although many TIL are bystanders, tumor-reactive clonotypes can be identified within TIL populations, and these cells bear transcriptional evidence of antigen encounter in the tumor microenvironment [37,38]. This is the conceptual distinction: vaccination begins with an antigenic instruction and asks whether the immune system can generate useful recognition; engagement-based approaches begin with biological evidence that recognition has already occurred and then seek to recover, enrich, and reinforce those cells. The strongest therapeutic version of this strategy is therefore not cell recovery alone or vaccination alone, but a coupled system in which a patient-specific antigenic representation is used first to recover tumor-relevant clonotypes and then to reinforce those same specificities after transfer.

### Rationale for reanalysis

Here, we reanalyzed and synthesized published tumor and peripheral-blood datasets in which tumor-reactive T-cell clonotypes were linked to physical interaction, antigen-dependent recovery, TCR sequence, transcriptional state, or functional validation. Rather than treating these studies as isolated observations, we evaluated whether they converge on a shared biological pattern: tumor-reactive T cells are enriched by productive engagement and preferentially occupy antigen-experienced states with preserved developmental potential.

### Tumor-reactive T cells must be selected, not merely nominated

Despite the known presence of antitumor T cells within TIL, their role as mediators of responses following immune checkpoint blockade, and the existence of candidate markers associated with them, the structure of the problem has obstructed many potentially scalable solutions. True tumor-reactive clonotypes are rare in most common cancers and are admixed with abundant bystander lymphocytes. They are also typically directed against mutation-derived antigens that are largely private to each patient. Sequencing and computation can narrow the candidate antigen space, but they cannot determine which antigens are naturally processed and presented by tumor cells, nor which of those are recognized by T cells present within the patient’s immune repertoire. Even increasingly sophisticated TCR–pMHC models cannot by themselves establish whether a living T cell can form productive contacts, integrate costimulatory and inhibitory signals, persist, proliferate, and generate useful effector progeny in vivo [39–42].

Approaches based on checkpoint-marker expression, cytokine release, peptide–MHC multimers, immunopeptidomics, and CRISPR-based knockout screens have been useful enrichment tools, but they remain surrogates rather than direct evidence of productive tumor recognition. Tellingly, the assays that come closest to ground truth are the least scalable. The clearest example is the current effort to “de-orphanize” tumor-reactive T cells within TIL, which typically requires testing against synthetic peptides or minigene constructs and often T-cell receptor cloning. Although new approaches may accelerate this work, de-orphanization remains bespoke, slow, and expensive [43].

Yet its benefits are real: de-orphanization can assign cognate antigen specificity, generate paired TCR–antigen datasets, guide vaccine reinforcement, and provide high-confidence labels for computational models. But these advantages do not remove the deeper problem. De-orphanization usually reconstructs specificity after the fact; it does not by itself recover the viable, developmentally useful T cells that have already converted tumor recognition into productive biological interaction. For a therapeutic platform, this distinction matters because bespoke reconstruction converts each patient into a slow target-discovery project, whereas recovery of viable engaged cells turns the patient’s own immune history into the starting material.

If current methods are limited by reliance on surrogates or laborious reconstruction, then a more direct alternative is to ask what scars tumor-antigen encounter leaves behind on T cells in vivo. This more direct strategy begins with first principles. When antitumor T cells recognize antigen in vivo, they do not simply upregulate a marker or release a cytokine. They enter a biological program marked by activation, proliferation, and extensive chromatin remodeling, with corresponding changes in gene expression, differentiation, and function. This program can include induction of inhibitory pathways and altered or delayed acquisition of cytotoxicity. Checkpoint signaling can dampen costimulatory and TCR-mediated signaling, limiting effector output without necessarily preventing antigen recognition or physical engagement with tumor cells or antigen-presenting cells, even when cytokine production or cytolytic activity is reduced [44–48]. These features suggest a different class of enrichment metrics for tumor-reactive T-cell discovery.

Single-cell studies have now made this state-based trace of tumor recognition visible. In melanoma, Oliveira et al. linked TCR specificity and molecular phenotype at single-cell resolution, showing that bona fide antitumor CD8+ T cells can be distinguished from bystander T cells by recurring phenotypic and transcriptional features [37]. Lowery et al. extended this logic across metastatic human cancers by mapping functionally validated neoantigen-reactive TCR clonotypes onto single-cell transcriptomes and deriving NeoTCR signatures that could prospectively enrich tumor-reactive receptors [49]. Caushi et al. used functionally identified mutation-associated neoantigen-specific TCRs as molecular barcodes to localize bona fide tumor-reactive clones within anti-PD-1-treated lung cancers [47], while Zheng et al. found that neoantigen-reactive T cells in gastrointestinal cancers preferentially occupied exhausted, CXCL13-enriched transcriptional states distinct from bystanders [50].

One of the most important insights from this work is that the imprint of tumor-antigen encounter can persist beyond the tumor microenvironment itself. Yossef et al. demonstrated that rare neoantigen-specific peripheral blood lymphocytes, termed NeoPBL, are detectable in the circulation, and that many of these validated antitumor clonotypes overlap with TIL. Compared with their intratumoral counterparts, circulating NeoPBL exhibit a less dysfunctional, memory-like transcriptional state, yet retain canonical features of antigen experience, including CD103, TIGIT, PD-1, and CD39 expression [38]. This blood-accessible signature creates a practical opportunity to enrich for and recover antitumor TCRs without relying exclusively on tumor-derived material.

More recent computational approaches extend this logic from descriptive signatures toward therapeutic prioritization. Tan et al. trained predicTCR using high-throughput TCR cloning and experimental reactivity validation, and reported improved prediction of tumor-reactive TCRs over earlier gene-set approaches [42]. Pétremand et al. similarly combined TRTpred with an avidity predictor to prioritize clinically relevant TCRs for personalized T-cell therapy [41].

These studies are important because they move beyond marker lists toward experimentally calibrated receptor prioritization. But they also illustrate the remaining constraint. Even high-quality tumor de-orphanization datasets, including those from Rosenberg and colleagues [38,49], remain modest and uneven relative to the diversity of TCR–pMHC recognition; newer screening efforts increase scale, but often by testing larger candidate-receptor sets under defined assay conditions rather than by broadly sampling HLA alleles, tumor types, antigen classes, processing contexts, and patient states. As a result, these approaches are best understood as powerful prioritization and reconstruction strategies: they nominate receptors or rank clonotypes for synthesis, cloning, expression, and validation, but they do not directly recover viable antigen-responsive cells that have already demonstrated productive tumor-relevant engagement.

Together, these studies establish an important principle: tumor recognition leaves a reproducible transcriptional imprint. But the data also caution against treating tumor reactivity as a single universal transcriptional state. CXCL13+CD39+PDCD1+ tumor-resident T cells in gastrointestinal cancer, mutation-associated neoantigen-specific TIL after PD-1 blockade in lung cancer, antitumor CD8+ T cells in melanoma, and circulating NeoPBL states are overlapping but not interchangeable. Shared features recur — antigen experience, checkpoint expression, tissue adaptation, CXCL13 or ENTPD1/CD39 expression in some tumor-resident populations, and preservation of progenitor or memory-like programs in other compartments — yet the exact signature varies with tumor histology, anatomical compartment, treatment exposure, antigenic context, timing, and patient-specific biology. That variation is likely biological, not merely technical noise. Thus, transcriptional state is best understood as a calibrated footprint of prior tumor-antigen encounter, not as tumor recognition itself [37,38,42,47,49–51]. Its strongest use is to prioritize candidate tumor-reactive clonotypes when anchored to TCR sequence, antigen validation, tumor recognition, or clinical function [52–54].

Cognate recognition tends to produce stable, long-lived cellular interactions, whereas nonspecific contacts are typically brief and transient [55–58]. General synapse studies establish that cognate recognition can stabilize T-cell contacts with antigen-presenting targets, while tumor-focused studies show that physically clustered or doublet-forming T cells can be enriched for tumor-reactive activity [59,60]. T-cell adherence to a tumor cell, or to an antigen-presenting cell displaying cognate peptide–MHC, therefore provides a physically observable trace of antigen recognition.

### Engagement is the footprint of recognition

The preceding studies define both the value and the limit of transcriptional inference: signatures can prioritize likely tumor-reactive clonotypes, but they do not by themselves establish native tumor recognition or deliver a viable therapeutic input population. Four practical gaps follow: antigen certainty, context dependence, therapeutic recoverability, and population loss during marker-based selection. First, transcriptional signatures report the consequences of prior stimulation; they do not prove the identity, tumor origin, or native presentation of the antigen recognized. A cell may look antigen-experienced, exhausted, tissue-resident, or recently activated, yet its TCR may recognize a bystander antigen, an assay-format peptide, or a target that is not naturally processed and displayed by tumor cells at sufficient density. This problem is illustrated by peptide-vaccine studies in which vaccine-elicited T cells recognized exogenous peptide but showed limited recognition of naturally processed tumor antigen or failed to prevent tumor progression despite large measurable antigen-specific responses [34,36].

Second, tumor-reactive signatures are context dependent. Melanoma antitumor CD8+ T cells, mutation-associated neoantigen-specific TIL after PD-1 blockade, CXCL13/ENTPD1-rich gastrointestinal tumor T cells, and circulating NeoPBL cells all carry evidence of tumor-antigen encounter, but they differ by tissue compartment, treatment exposure, differentiation state, cytotoxic program, and degree of progenitor or memory-like preservation [37,38,42,47,50].

Third, signature-based discovery often produces a candidate receptor, not a therapeutic cell. If the relevant cell has been consumed by single-cell sequencing, the investigator has a TCR sequence that must still be synthesized or cloned, expressed, validated, and assessed for safety and utility [41,42,49].

Fourth, marker-based recovery is necessarily lossy. PD-1, CD39, CXCL13, CD103, 4-1BB, OX40, or combinations of these markers can enrich tumor-reactive cells, but each gate also narrows the recovered population and risks excluding bona fide tumor-reactive cells that fall outside the expected state because of histology, compartment, treatment, antigen dose, timing, or patient-specific biology [52–54]. Thus, transcriptional signatures are powerful evidence that recurring tumor-reactive states exist, but they remain tools for prioritization. They do not replace a functional enrichment principle that recovers viable cells after productive antigen-dependent interaction.

Engagement captures a proximal physical event between TCR binding and downstream effector function. Here, engagement means stable, antigen-dependent physical interaction between a T cell and a tumor cell or antigen-presenting cell, including synapse formation, adhesion, and sustained cell–cell contact [55–60]. It is not identical to TCR binding, and it is not equivalent to cytokine release, proliferation, cytolysis, or transcriptional remodeling. These layers can be uncoupled: a tumor-reactive T cell may recognize and adhere to its target while showing limited cytotoxicity or attenuated cytokine production, whereas activation markers or cytotoxic programs may identify antigen-experienced cells without proving tumor specificity.

Checkpoint signaling makes this separation especially important. In chronically stimulated tumors, inhibitory receptors can dampen TCR signaling and suppress immediate effector output without necessarily preventing stable contact with antigen-presenting cells or tumor cells [44–46,48,61,62]. Thus, even when cytotoxicity is muted, tumor-reactive T cells may still retain the capacity to form biologically meaningful conjugates [55,56,59]. Engagement can therefore reveal clonotypes that phenotype-only or function-only assays may miss.

Across the settings considered here, engagement does not converge on a single canonical phenotype, but it repeatedly points to a shared biological theme: antigen experience coupled to preserved developmental potential. Engaged cells are enriched within antigen-experienced compartments that overlap with progenitor-associated programs rather than terminal effector differentiation alone [37,44,45,59,63–65]. This suggests that tumor-reactive T cells may preferentially reside in a conserved developmental reservoir: cells that have seen antigen, retained proliferative potential, and remained poised for reinvigoration. Engagement is therefore not merely a technical readout of cell contact. It is a way to recover the physical history of tumor recognition from within a complex and misleading immune landscape.

### Reanalysis of published datasets

We focused on studies in which tumor-reactive T cells could be evaluated across at least one of four dimensions: physical engagement with tumor cells or antigen-presenting cells, validated neoantigen or tumor recognition, overlap between blood and tumor repertoires, and transcriptional evidence of progenitor-like versus terminal cytotoxic differentiation. The following sections organize these datasets by anatomical source and experimental logic.

### Within tumors, physical engagement exposes the reactive compartment

Ibáñez-Molero and colleagues identified heterotypic clusters of CD8+ T cells physically interacting with tumor cells, antigen-presenting cells, or both across 21 melanoma metastases [59]. Single-cell RNA and TCR sequencing demonstrated that these clustered T cells were clonally expanded and enriched for tumor-reactive transcriptional programs relative to singlet T cells, which were dominated by bystander virus-reactive populations. Following expansion, cluster-derived T cells exhibited markedly increased cytokine production and approximately nine-fold greater tumor-killing activity ex vivo, as well as superior tumor control in vivo. Notably, standard single-cell workflows typically exclude such clusters as presumed doublets, indicating that conventional analyses systematically discard a functionally important compartment of tumor-reactive T cells.

The phenotypic structure of these clusters was highly informative. Replotting the cell-state enrichment analysis as effect sizes emphasizes that, relative to T-cell singlets, tumor-associated T-cell clusters were depleted for naïve, memory, early effector-memory, and effector-memory states, but strongly enriched for a TCF7+ stem-like exhausted state **(Fig. 1).** Thus, physical clustering does not simply recover a broad mixture of activated or differentiated T cells. It preferentially enriches for a restricted antigen-experienced compartment with stem-like, persistence-associated features.

**Figure 1.**
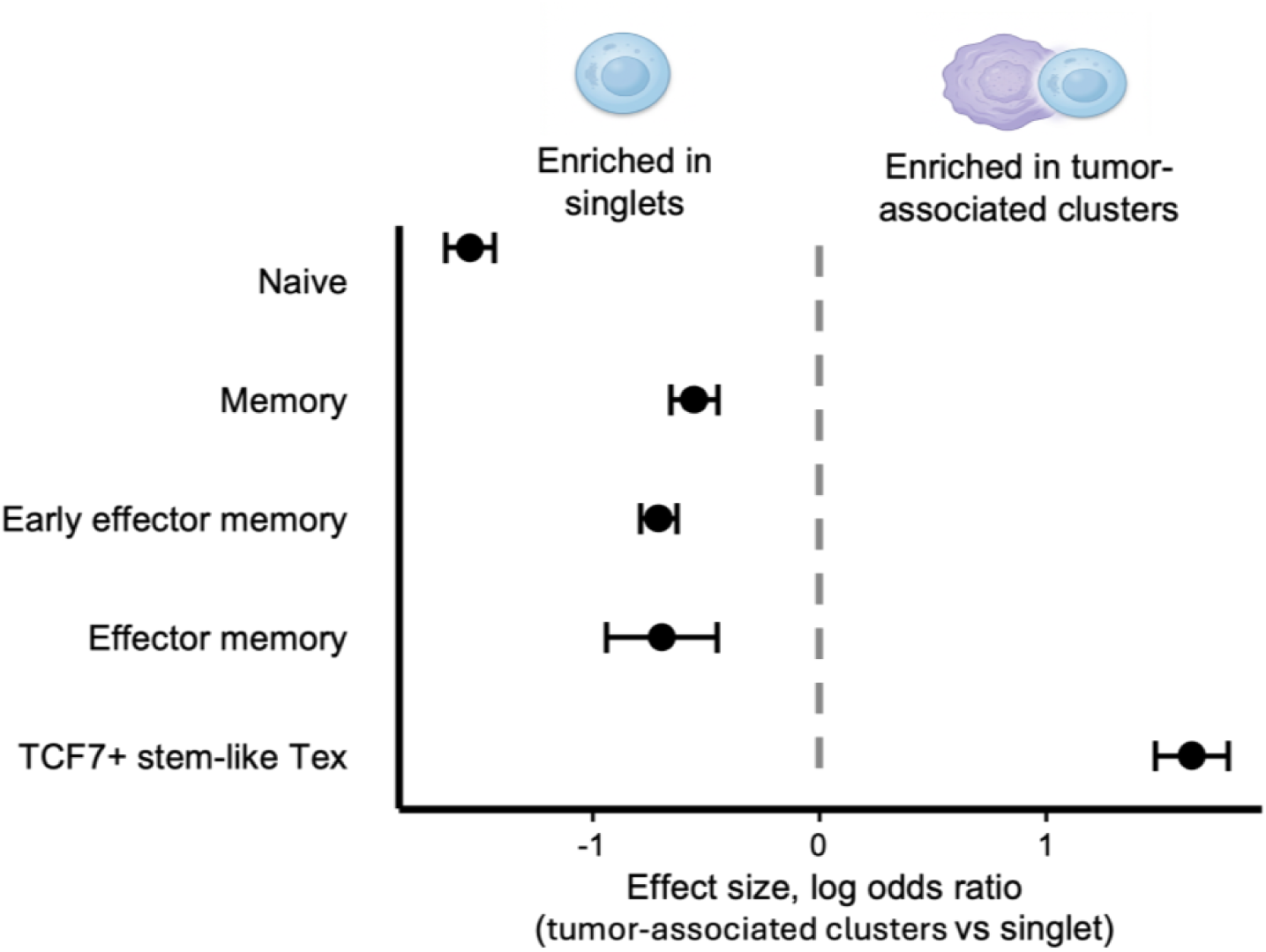
Tumor-associated T-cell clusters are enriched for a TCF7+ stem-like exhausted phenotype. Replotted log odds ratios compare CD8+ T cells from tumor-associated heterotypic clusters with CD8+ T-cell singlets. Values were replotted from the published cell-state enrichment analysis. Tumor-associated clusters were depleted for naive, memory, early effector-memory, and effector-memory states, but strongly enriched for a TCF7+ stem-like exhausted phenotype. Thus, clustering preferentially enriches an antigen-experienced progenitor-like compartment.

The lack of enrichment for effector-memory states is notable but biologically consistent with the nature of the assay [59]. Stable heterotypic clusters likely capture cells capable of sustained antigen-dependent signaling and target adherence, rather than cells already biased toward rapid cytotoxic execution [59,60]. Cells committed to cytolytic execution may undergo synaptic remodeling, centrosome and granule polarization, cortical actin clearance at the secretory domain, and eventual target disengagement [57,66–69]; such behavior could make terminally cytotoxic cells less likely to be recovered as persistent tumor-cell or APC-associated clusters in a static single-cell snapshot. Thus, the absence of effector-memory enrichment should not be interpreted as absence of tumor reactivity [52,53,59]. Rather, it supports the idea that productive adherence preferentially enriches for an antigen-experienced progenitor-like reservoir from which downstream cytotoxic progeny can arise [44,46,65].

Together with the functional data, this enrichment for TCF7+ stem-like exhausted cells links physical clustering to a progenitor-like population associated with clonal persistence, therapeutic response, and downstream effector potential. Antigen engagement thus places enriched tumor-reactive activity within a conserved antigen-experienced progenitor compartment outside the classical naïve–effector continuum. We next asked whether this same compartment could be detected among circulating tumor-reactive T cells.

### In blood, rare neoantigen-reactive cells converge on the same state

A complementary view comes from the blood. In an independent peripheral blood dataset from patients whose TIL had already been shown to contain neoantigen-reactive clonotypes, functional screening coupled to single-cell RNA and TCR profiling identified 36 validated neoantigen-specific clonotypes across six patients with metastatic solid tumors [38]. Set against a vast background of unrelated circulating T cells, including antiviral populations, these rare tumor-reactive clonotypes did not disperse randomly through transcriptional space. Instead, they converged in a restricted cluster, designated C9, that was distinct from bystander virus-reactive populations across all patients examined.

This circulating compartment closely mirrored the state emerging from tumor-based analyses. Neoantigen-reactive cells (NeoPBL) express progenitor-associated genes such as TCF7 in the setting of checkpoint expression, while showing little evidence of full cytotoxic differentiation. They stand in sharp contrast to EBV- and CMV-reactive populations, which occupy a more classically effector transcriptional space, with lower expression of progenitor-associated genes and higher expression of cytotoxic programs **(Fig. 2)**. That contrast is informative. Acute antigenic stimulation typically drives loss of stemness-associated genes together with acquisition of effector genes [70–76].

**Figure 2.**
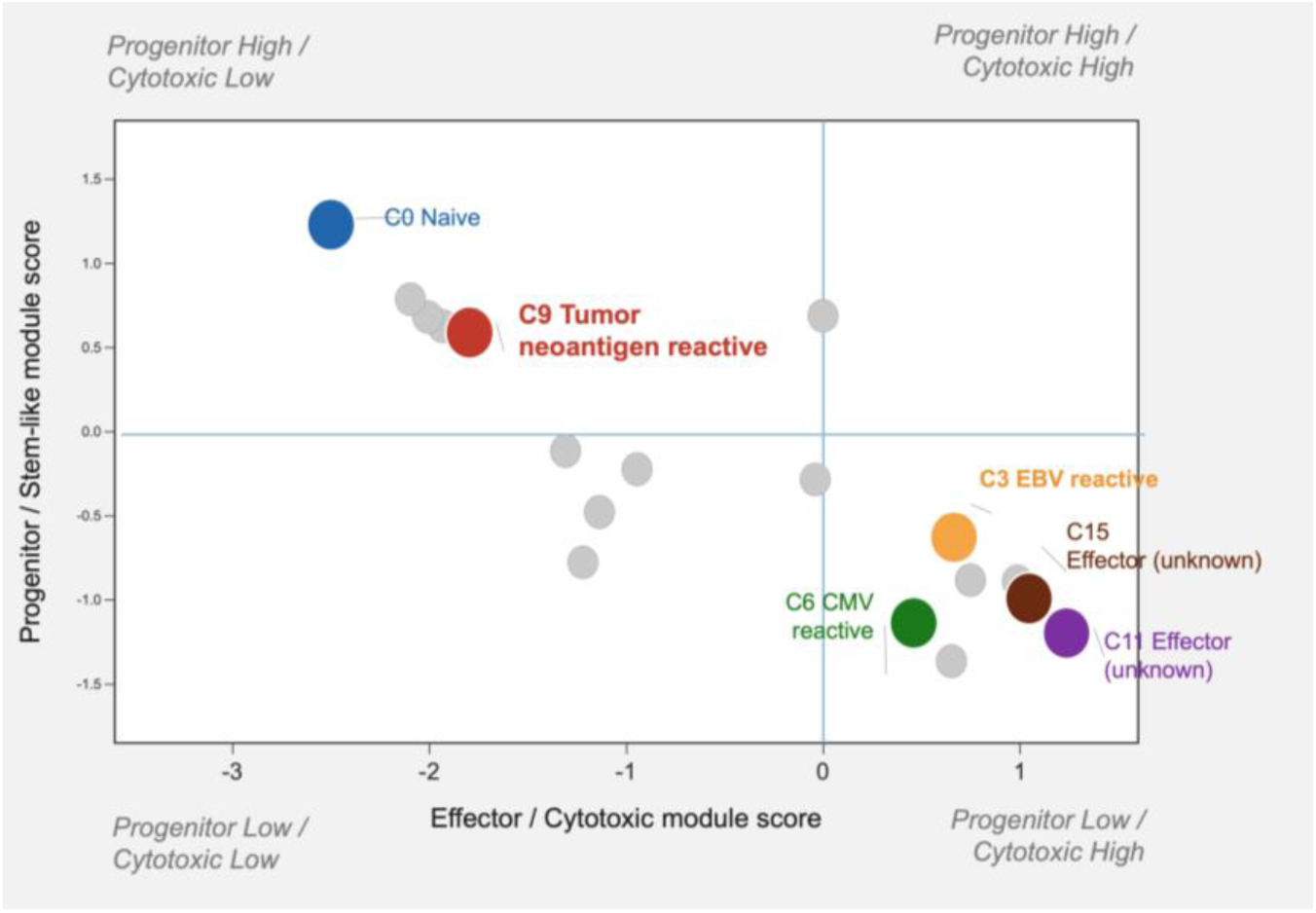
Circulating neoantigen-reactive CD8+ T cells occupy a progenitor-high, cytotoxic-low transcriptional state. Module-score analysis of peripheral blood CD8+ T-cell transcriptional clusters, plotted by progenitor/stem-like score versus effector/cytotoxic score. Scores represent mean log2 fold-change values for reported cluster-defining genes overlapping canonical progenitor/stem-like and cytotoxic-effector gene sets. The tumor neoantigen-reactive cluster C9 localizes to a progenitor-high, cytotoxic-low region, distinct from EBV- and CMV-reactive clusters, which occupy more cytotoxic effector-biased states.

Circulating tumor-reactive T cells did not fit either end of a simple naïve-to-effector progression. Rather, NeoPBL combined progenitor/trafficking-associated transcripts, activation and checkpoint-associated molecules, and stress-adaptation programs, while lacking the canonical cytotoxic-effector genes that dominate CMV- and EBV-reactive populations [38] **(Fig. 3)**. Thus, circulating tumor-reactive CD8+ T cells are not simply weak effectors; they occupy a distinct developmental compartment: antigen-experienced, chronically engaged, progenitor-skewed, and not yet terminally cytotoxic [77,78].

**Figure 3.**
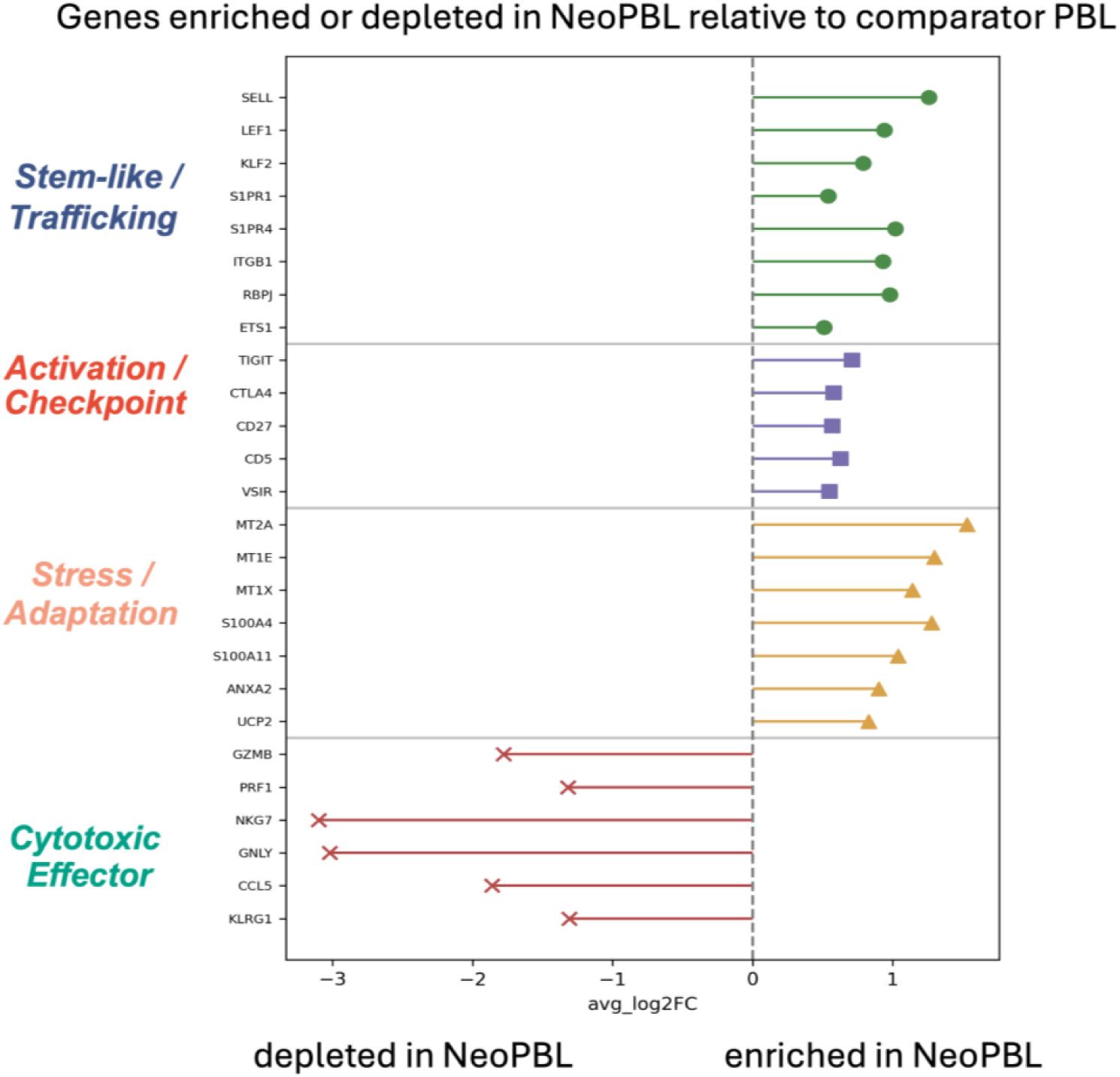
NeoPBL cells combine progenitor trafficking, checkpoint activation, and stress-adaptation programs while lacking cytotoxic-effector differentiation. Differential gene-expression structure of circulating neoantigen-reactive CD8+ T cells. Gene programs are summarized from reported differential-expression patterns. NeoPBL are enriched for progenitor/trafficking, activation/checkpoint, and stress-adaptation programs. In contrast, canonical cytotoxic-effector genes are depleted. This composite pattern defines a restrained tumor-reactive state: antigen-experienced, checkpoint-positive, developmentally restrained, and not terminally cytotoxic.

This point reframes the question. If tumor recognition is linked to this progenitor-like chronic-response state, then tumor-reactive clonotypes should not only map to a transcriptomic state associated with both stemness and checkpoint expression, with low cytotoxic-effector gene expression – they should preferentially expand within it.

### Expansion occurs within the progenitor-like reservoir, not outside it

Neoantigen-reactive clonotypes circulating in blood can be highly expanded, in some cases reaching clone sizes comparable to viral-reactive clonotypes. However, these cells remain transcriptionally distinct from classical antiviral effector populations. Clonotypes of unknown specificity are broadly distributed across transcriptional space but are rarely both highly expanded and strongly aligned with the NeoPBL program. Thus, chronic tumor-antigen exposure appears compatible with clonal expansion while constraining terminal cytotoxic differentiation **(Fig. 4)**.

**Figure 4.**
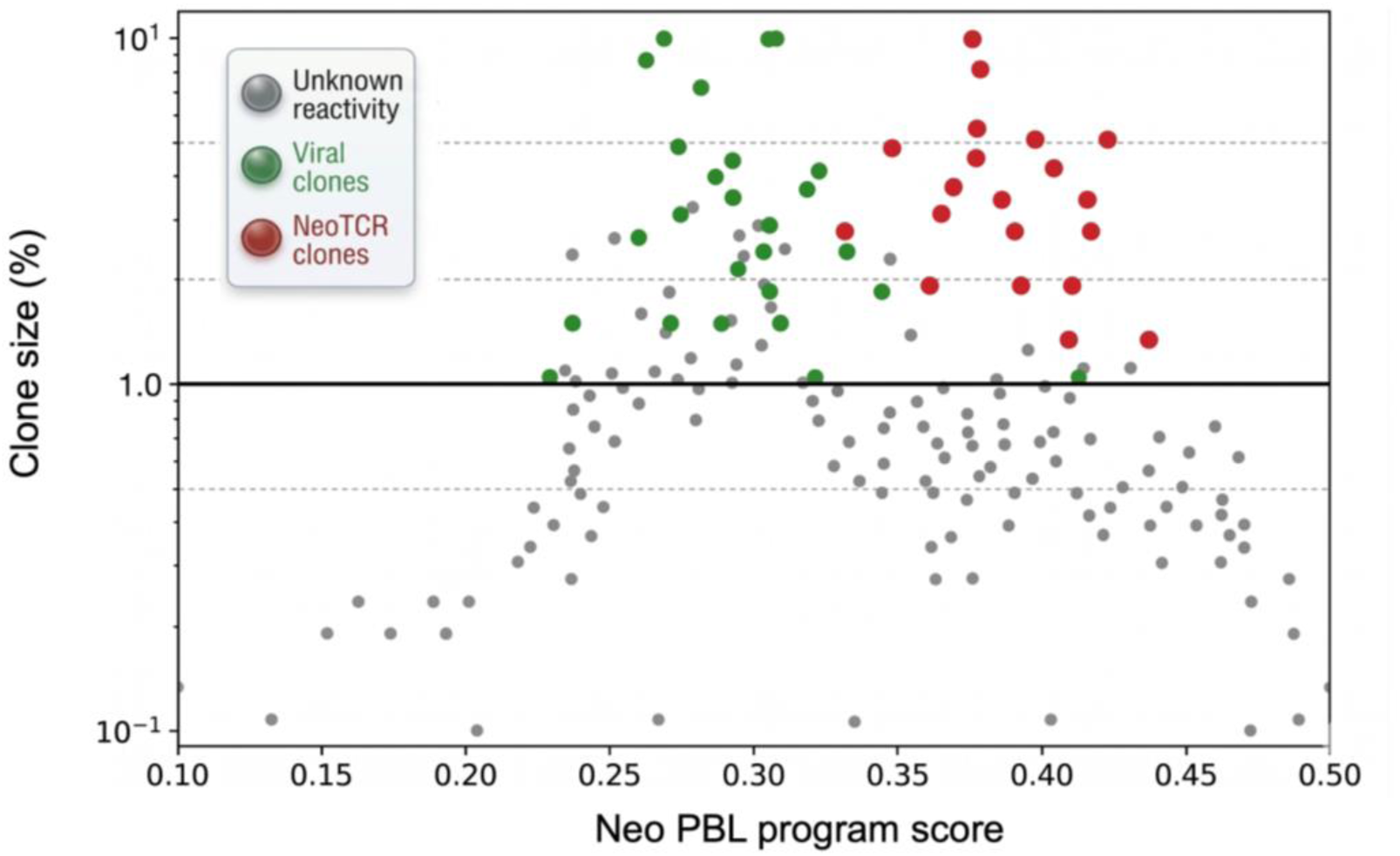
Expanded neoantigen-reactive clonotypes are concentrated within the NeoTCRPBL gene-expression program. Relationship between clonotype size and NeoTCRPBL gene-set enrichment score among circulating CD8+ T-cell clonotypes. Clonotype-size and program-score relationships were reorganized from the published NeoTCRPBL analysis. Neoantigen-reactive clonotypes occupy a high NeoTCRPBL program-score region and include clonotypes expanded to frequencies comparable to antiviral populations. Viral-reactive and unknown-specificity clonotypes are more broadly distributed, whereas unknown clonotypes are rarely both highly expanded and strongly aligned with the NeoTCRPBL program.

Within this framework, progenitor-exhausted CD8+ T cells (TPEX) provide a useful reference point for interpreting the transcriptional state observed here. TPEX cells arise under conditions of persistent antigen exposure and are characterized by co-expression of inhibitory receptors together with stem-associated transcription factors such as TCF7 (TCF-1) and LEF1, which preserve developmental potential within the CD8+ lineage. These cells are antigen-experienced yet remain distinct from terminally exhausted or fully cytotoxic effector populations, retaining proliferative capacity and the ability to generate downstream effector progeny [44,45,61,62].

However, the datasets considered here suggest that tumor-reactive T cells should not be defined narrowly by a single transcriptional identity such as TPEX [52,53,65]. Rather, tumor-reactive T cells occupy a broader antigen-experienced state that emerges through repeated encounter with tumor-derived antigen across linked lymphoid and tumor compartments [52,53,79–81]. Cross-presenting dendritic cells in tumor-draining lymph nodes likely initiate priming and help maintain a progenitor-like reservoir, whereas subsequent encounter with malignant cells and intratumoral antigen-presenting cells can reinforce antigen experience, checkpoint expression, and partial developmental arrest [80–84]. Within this state, cells retain progenitor-associated transcriptional programs while cytotoxic differentiation remains constrained [44,46,77,80,81]. Inhibitory receptor expression within this compartment likely reflects regulatory signaling imposed by persistent antigen exposure rather than irreversible commitment to terminal exhaustion [52–54,61,62].

Consistent with this interpretation, checkpoint blockade appears to act preferentially on TCF7+ progenitor-like T cells, which proliferate and generate cytotoxic progeny after PD-1 pathway inhibition [44,45,64]. This activity need not occur only at a direct tumor-cell/T-cell interface; APC-rich niches in tumors and tumor-draining lymph nodes may be critical sites of PD-1–PD-L1 regulation. In several models, PD-L1 expression by host myeloid cells or dendritic cells regulates checkpoint response, and PD-L1 blockade in tumor-draining lymph nodes can seed tumors with progenitor-exhausted CD8+ T cells [85–87]. The same principle applies to adoptive immunotherapy, where products containing progenitor-like cells appear more effective than products dominated by terminally differentiated cells [65,76].

### Engagement-based capture recovers circulating tumor-reactive T-cell clonotypes

The tumor-reactive T cells most relevant to durable tumor destruction appear to be enriched within a recurrent antigen-experienced, progenitor-like compartment. The practical question, and the one that matters for a therapeutic platform, is whether this compartment can be recovered prospectively from accessible patient material. Blood recovery is not just biologically interesting; it changes logistics, repeatability, and tumor-type reach. If this compartment retains the capacity to convert antigen recognition into productive cellular interaction, then engagement can be used as a prospective enrichment variable in more than one format: direct physical adherence in conjugate-based assays, recovery of APC–T-cell conjugates after cognate antigen encounter, or short-term assays that link antigen-triggered activation to recovery of viable cells.

We therefore examined the study by Khateb et al., which tested whether rare neoantigen-reactive peripheral blood CD8+ T cells could be enriched using NeoSelect, a strategy combining neoantigen stimulation with bead extraction of peptide-pulsed target cells [63]. In this approach, autologous CD4+ T cells expressing truncated NGFR are pulsed with patient-specific minimal neoantigen peptides and used as target APCs. After short-term co-culture, NGFR+ target cells are bead-extracted, enriching antigen-responsive CD8+ T cells associated with the peptide-pulsed targets. Thus, the method is best described as antigen-specific target-cell extraction coupled to expansion, rather than receptor prediction, peptide–MHC modeling, or direct sorting of activation-marker-positive cells.

An earlier AML proof-of-principle deserves mention as a prescient but lower-resolution antecedent [60]. By physically sorting stable T cell–tumor doublets, that study attempted to use engagement itself as the enrichment variable and recovered a fraction with greater cytotoxic activity than the cells left behind. But the work remained phenotypically shallow and could not determine whether doublet formation isolated bona fide tumor-reactive clonotypes or more broadly selected activated, adhesion-prone effector cells. Khateb et al. place the same underlying principle on firmer ground by linking engagement-based recovery to TCR sequence, neoantigen specificity, tumor overlap, and single-cell transcriptional state [63].

We then asked whether clonotypes recovered from blood by this higher-resolution engagement-based strategy corresponded to tumor-reactive populations observed in vivo. TCR sequences from NeoSelect-expanded blood populations were compared with single-cell TIL atlases from matched tumors [63]. Many of the recovered blood-derived clonotypes were also detected within the tumor T-cell repertoire, where they localized predominantly to exhausted/dysfunctional neoTCR8-like compartments expressing markers such as CXCL13, ENTPD1, PDCD1, and TIGIT. At the same time, the expanded circulating NeoPBL retained a distinct ex vivo program enriched for memory and costimulatory features, including IL7R, TCF7, KLF2, CD28, and CD40LG. This pairing is the key biological point: NeoSelect can recover rare circulating counterparts of tumor-experienced TIL clonotypes while yielding a less terminally differentiated ex vivo transcriptional state. Across patients and neoepitopes, the method rescued clonotypes that were undetectable or present at extremely low frequency in baseline blood and expanded them into dominant populations after in vitro culture. These findings support the idea that engagement-based recovery from blood can prospectively enrich for physiologically relevant candidate tumor-reactive clonotypes, which can then be linked to tumor overlap, neoantigen specificity, TCR sequence, and transcriptional state.

### Engagement-recovered NeoPBL are developmentally restrained, not inert

The circulating neoantigen-reactive CD8+ cells recovered by engagement are not classical effectors. They carry a mixed program of antigen experience, developmental reserve, and retained costimulatory or adhesive competence: TCF7, IL7R, KLF2, and SELL; CD27 and CD28 alongside PDCD1, TIGIT, TOX, IL2RA, and CD40LG. Relative to reactive TIL, NeoPBL skew toward IL2 and IL21 rather than IFNG and TNF.

CD226 (DNAM-1) is particularly informative, because CD226-dependent costimulation has been linked to effective anti-PD-(L)1 responses and is differentially expressed on TIL [88]. The coexistence of CD226 with TIGIT is notable because these receptors represent opposing functional arms of the PVR/nectin ligand axis: CD226 provides activating costimulation through shared ligands such as CD155/PVR and CD112/PVRL2, whereas TIGIT is an inhibitory receptor that can limit CD226-dependent signaling [89]. Thus, checkpoint expression in NeoPBL should not be read simply as terminal exhaustion or irreversible dysfunction; it coexists with preserved adhesive and costimulatory circuitry, placing these cells closer to a restrained progenitor-like state than to a classical cytotoxic effector state: chronically engaged, developmentally restrained, and not yet terminally cytotoxic **(Fig. 5)**. Together, the summary program view and direct pairwise comparisons distinguish NeoPBL from both non-reactive circulating cells and more cytotoxic tumor-resident reactive TIL **(Figs. 3 and 5)**.

**Figure 5.**
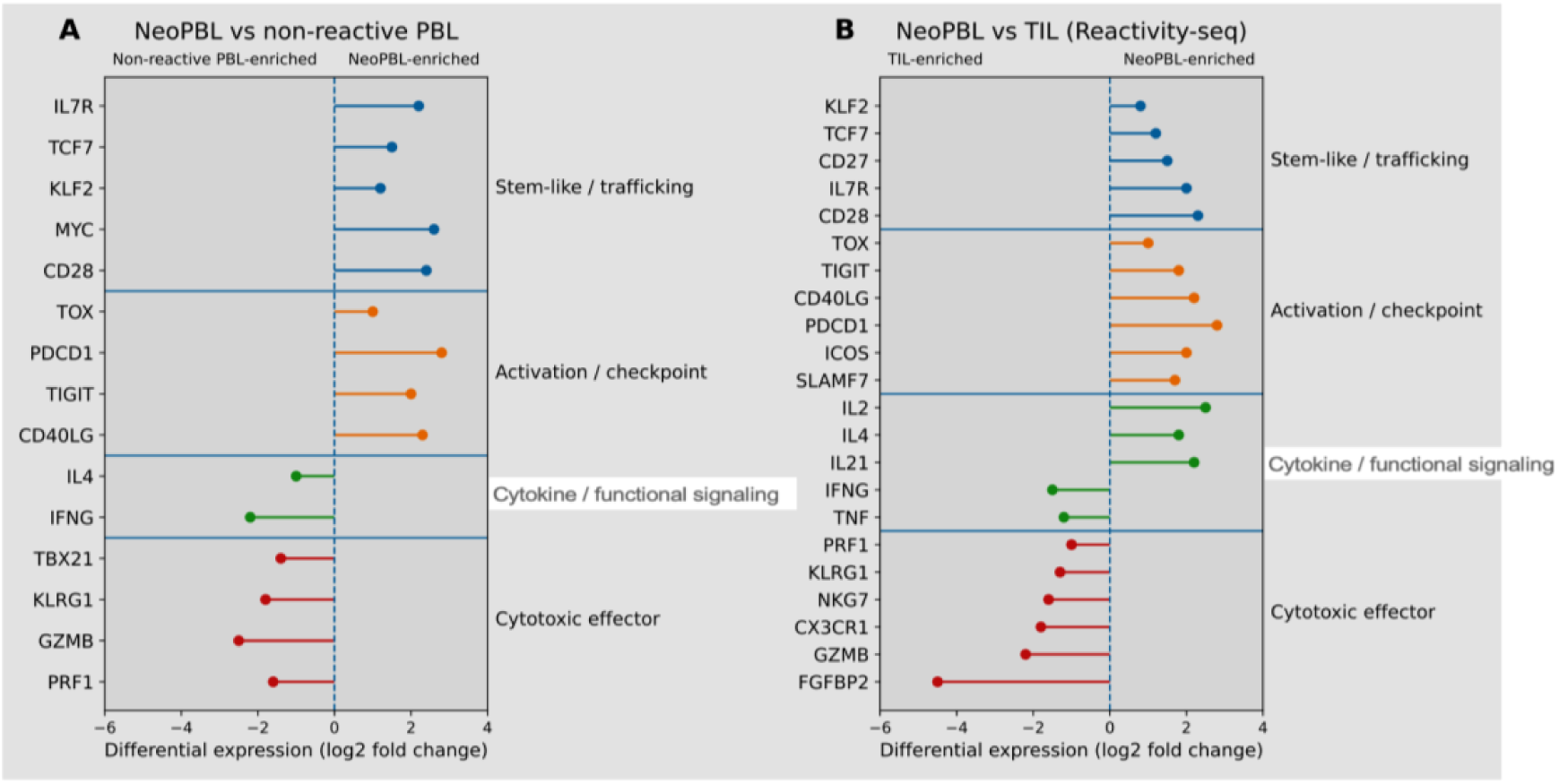
Circulating neoantigen-reactive CD8+ T cells display a restrained progenitor-like program distinct from both non-reactive blood T cells and more cytotoxic reactive TIL. Differential gene-expression analysis of circulating neoantigen-reactive peripheral blood lymphocytes (NeoPBL). Pairwise comparisons summarize differential-expression relationships between NeoPBL, non-reactive PBL, and reactive TIL. A, compared with non-reactive peripheral blood lymphocytes, NeoPBL are enriched for stem-like/trafficking and activation/checkpoint programs, while cytotoxic-effector genes are relatively depleted. B, compared with neoantigen-reactive TIL identified by Reactivity-seq, NeoPBL retain higher progenitor/trafficking, costimulatory, and selected cytokine programs, with lower cytotoxic-effector expression. Together, these data define circulating tumor-reactive T cells as antigen-experienced and checkpoint-positive but developmentally restrained and not terminally cytotoxic.

Figure 6 summarizes this therapeutic architecture as a three-step workflow: engagement-based recovery defines the selected input population, stemness-preserving expansion amplifies the recovered clonotypes while maintaining developmental reserve, and targeted deployment pairs adoptive transfer with in vivo antigenic reinforcement to support localized expansion and persistence. Taken together, these observations support a unifying principle: physical engagement enriches for tumor-reactive T cells, and these cells are preferentially maintained in an antigen-experienced developmental reservoir that sustains effective antitumor immunity.

**Figure 6.**
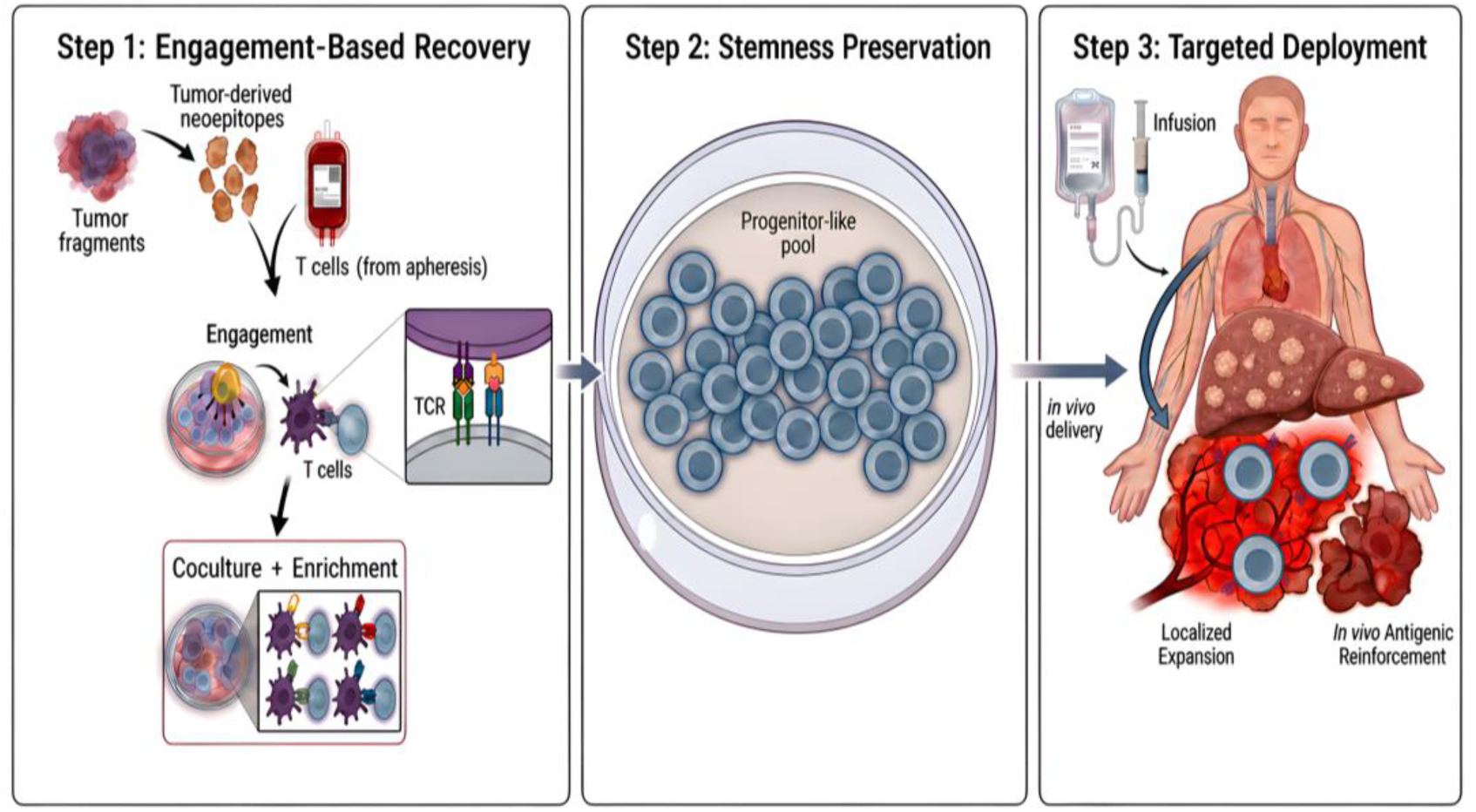
Three-module therapeutic architecture for recovery, expansion, and reinforcement of tumor-reactive T cells. Three-step therapeutic architecture for engagement-based recovery, stemness preservation, and targeted deployment of tumor-reactive T cells. Step 1, engagement-based recovery, uses patient-specific tumor-derived neoepitopes, tumor material, and peripheral T cells to enrich viable T cells that form productive antigen-dependent interactions with tumor cells or autologous antigen-presenting cells. Step 2, stemness preservation, expands the recovered clonotypes under conditions intended to maintain a progenitor-like pool rather than drive terminal cytotoxic differentiation. Step 3, targeted deployment, returns the selected cells to the patient and pairs adoptive transfer with in vivo antigenic reinforcement to support localized expansion and persistence at tumor sites. Together, the workflow links empirical recovery of tumor-engaged T cells, preservation of developmental reserve, and antigen-matched reinforcement after transfer.

### Convergent biological themes

Across these reanalyses, the individual datasets differ in tumor type, compartment, assay format, and validation strategy. Nevertheless, they point to a recurring biological architecture: productive engagement enriches tumor-relevant clonotypes, and these clonotypes are often found in antigen-experienced states that retain proliferative and developmental reserve. The following sections interpret that convergence and its implications for therapeutic cell recovery.

### Why adherence functionally discriminates tumor-reactive T-cell states

Productive target adherence provides a functional way to capture tumor relevance rather than infer it indirectly. By selecting cells that physically engage a cognate target, adhesion-based workflows enrich for T cells that have translated recognition into sustained cellular interaction, with preserved synapse competence, integrin coupling, and signal-integration capacity, while still requiring validation for antigen specificity and native tumor recognition. Stable adherence may also capture an earlier, signal-integrative phase of antigen engagement, before full cytolytic execution and target disengagement. Thus, productive adhesion provides a physical link between molecular specificity and recoverable cellular behavior. If engagement enriches for functionally relevant tumor-reactive T-cell states, the next question is where these cells reside in vivo.

### Tumor-reactive T cells reside beyond the tumor

The tumor is the most obvious place to look for tumor-reactive T cells, and the place where the field began [90], but it is not the only compartment in which such cells reside, and it may not be the best place to recover them. TIL remain the strongest clinical proof that a T cell has encountered cancer in vivo. But a tumor biopsy is a spatially and temporally biased snapshot: sampling is limited, infiltration is uneven, and many relevant clonotypes may be excluded from the tumor mass, trapped in stromal or vascular niches, retained in draining lymph nodes, or circulating in blood.

Tumor recognition is therefore not a local event confined to the lesion. It unfolds across a connected lymph node–blood–tumor axis. Tumor antigens are captured and cross-presented by dendritic cells in draining lymph nodes, reinforced by subsequent encounters with malignant cells or intratumoral APCs, and echoed systemically by clonally related T cells in blood. The tumor can show that recognition occurred, but the surrounding immune circuit may preserve the cells most useful for recovery and therapeutic amplification.

This distinction matters because intratumoral T cells are often the product of repeated antigen exposure. They acquire tissue-resident features, inhibitory receptors, and more differentiated or dysfunctional states. By contrast, clonally related tumor-reactive cells in lymphoid tissues and blood can be less terminally differentiated and more developmentally plastic. Across mouse models and human datasets, tumor-draining lymph nodes can preserve TCF1/TCF7+ tumor-specific CD8+ reservoirs that maintain antitumor immunity and seed more differentiated intratumoral progeny. Paired blood–tumor studies separately show that clonally related T cells can occupy divergent circulating and intratumoral states [80,81,84].

Thus, the TIL compartment should not be treated as the complete tumor-reactive repertoire. It is a biased enrichment for cells that have already endured persistent antigen exposure within the tumor microenvironment. The circulating and lymphoid compartments may instead preserve an earlier, more proliferative reservoir: rare, antigen-experienced clonotypes capable of sustaining long-term responses and replenishing the tumor.

This also changes how peripheral blood should be viewed. Tumor-specific T cells in blood are rare, and their frequency cannot be inferred simply from tumor mutational burden, predicted neoantigen load, or standard cytotoxic-response readouts. But rarity does not mean irrelevance. Blood may provide a window into the less exhausted, more renewable fraction of the antitumor response. Consistent with this interpretation, the magnitude of systemic neoantigen-specific CD8+ T-cell responses, rather than tumor mutational burden or predicted neoantigen load alone, has been associated with clinical outcome after PD-(L)1 blockade [91]. Peripheral blood is therefore not merely a convenient source of cells. It is a distinct biological compartment in which tumor recognition may remain visible before it is fully obscured by the selective pressures of the tumor microenvironment. This matters operationally because blood can be sampled repeatedly, processed earlier, and paired with tumor sequencing or antigen nomination without requiring that the final therapeutic product be derived entirely from a surgically accessible tumor lesion.

### Why selection must precede expansion

Expansion is often treated as a neutral manufacturing step that occurs after tumor-reactive T cells have been identified or obtained. But expansion is itself a selection event. Culture conditions impose their own selection pressures, favoring cells that survive antigen withdrawal, tolerate cytokine exposure, respond to nonspecific stimulation, and proliferate rapidly in vitro. If tumor-reactive clonotypes are rare, developmentally restrained, or partially inhibited by persistent antigen exposure, expansion-first workflows can amplify the wrong cells before the right ones have been identified. Recent work further suggests that even controlled T-cell/tumor interactions can remain probabilistic, with rare IFN-γ-poised T cells initiating paracrine amplification of MHC-I expression and tumor killing [92]. This reinforces the need to recover not only tumor-reactive clonotypes, but also functional states capable of propagating productive tumor recognition.

This is particularly important for bulk TIL manufacturing. In the expansion-first studies considered here, manufacturing relied on strong ex vivo growth and activation conditions, including IL-2, nonspecific TCR stimulation, feeder-supported expansion, and, in the Amaria trial, agonistic 4-1BB stimulation [51,93]. These conditions can generate large numbers of activated T cells, but they do not necessarily preserve the tumor-reactive fraction. Instead, they preferentially expand cells best adapted to the culture conditions provided. Bystander memory cells, tumor-irrelevant antigen-experienced cells, or clonotypes with greater intrinsic proliferative capacity may outgrow rarer tumor-reactive cells that have been shaped by chronic antigen exposure, checkpoint signaling, and tissue adaptation. In this setting, expansion does not merely fail to discriminate; it can actively reshape the product.

A longitudinal melanoma TIL-ACT study by Chiffelle et al. illustrates both the value and the interpretive risk of studying expansion-first products. By tracking TCR clonotypes from baseline tumors into ACT products and post-treatment samples, the authors showed that response was associated with pre-existing tumor-reactive CD8+ clonotypes in the baseline tumor, their effective mobilization into the expanded product, and their preferential re-entry into tumor after transfer. They also reported loss of tumor-associated transcriptional signatures, including exhaustion, during expansion, which they interpreted as reinvigoration or reprogramming. But clonotype tracking is not single-cell fate mapping. A shared TCR identifies a clonotype, not the fate of an individual exhausted cell. Apparent “reprogramming” at the clonotype level may therefore reflect selective outgrowth of pre-existing progenitor-like, less terminally exhausted, or otherwise culture-adapted variants within tumor-reactive clonotypes rather than durable erasure of an exhaustion-imprinted state. The data strongly support mobilization and selection of tumor-reactive clonotypes; they do not prove that exhausted TIL are durably reprogrammed into renewable therapeutic cells [51].

This distinction becomes even more consequential outside melanoma, where tumor-reactive TIL are often rarer, less abundant, and less competitively represented in bulk cultures. The Amaria phase II trial in colorectal, pancreatic, and ovarian cancer provides the clearest clinical example among the studies considered here. In that trial, manufacturing incorporated IL-2, anti-CD3 stimulation, agonistic 4-1BB stimulation, rapid expansion, lymphodepletion, and post-infusion high-dose IL-2 [93]. The process was technically feasible and generated large CD8-enriched, activated TIL products, but produced no objective responses among 16 treated patients. The disease-control signal was driven by stable disease, declined substantially by the 12-week primary efficacy time point, and was accompanied by a median progression-free survival of only 2.53 months. Thus, successful manufacturing of bulk TIL showed that cells could be grown, not that the therapeutically relevant clonotypes had been selected.

A similar caution applies to modern personalized neoantigen-specific cell products generated through peptide-driven stimulation. In the BNT221 phase 1 melanoma study, autologous peripheral-blood T cells were expanded against patient-specific neoantigen peptides and returned as a personalized polyclonal product. The study demonstrated feasibility of manufacturing blood-derived neoantigen-specific T-cell products, but in the completed monotherapy cohort only nine patients were treated; best overall response was stable disease in six patients, with tumor reductions of no more than 20% reported in four of those patients [94]. Thus, peptide-driven product generation can produce a plausible, mutation-reactive cellular therapy without establishing that the therapeutically decisive clonotypes have been selected, expanded, and positioned to mediate regression of established tumors.

Together, these examples sharpen the practical problem. The key question is not whether T cells can be expanded. They can, both in vivo through vaccination or ex vivo under highly artificial culture conditions. The question is whether the right clonotypes are present in the input population before expansion begins. For rare tumor-reactive T cells occupying restrained, antigen-experienced states, selection must precede amplification. Otherwise, manufacturing risks converting a biological discovery problem into a culture-competition problem, where the cells that grow best are mistaken for the cells that matter.

Across the datasets considered here, engagement repeatedly converges on a family of antigen-experienced states in which developmental reserve is preserved, and terminal cytotoxic differentiation is restrained. This biology explains why selection before expansion matters: the therapeutic objective is not merely to reactivate end-stage effectors, but to recover clonotypes capable of renewed proliferation, differentiation, and persistence. Engagement-based selection is useful precisely because it captures this recoverable state before it is lost to terminal differentiation or obscured by expansion of irrelevant lymphocytes.

### Boundaries of engagement-based selection

This framework has important boundaries. Physical adherence is not, by itself, proof of recognition of naturally processed tumor antigen on malignant cells in vivo. Even when engagement is antigen-dependent, its meaning depends on the identity of the ligand, the density of peptide–MHC, the presenting cell, and the tissue context [95–97]. T cells may engage peptide-loaded APCs or experimentally engineered antigen constructs yet still differ in their ability to recognize endogenously processed antigen displayed by tumor cells in vivo [95,96]. This is the same limitation faced by peptide-stimulation workflows: they can demonstrate recognition of the assay format without proving productive engagement of native tumor at physiologic antigen density [32–36]. Similarly, the link between restrained progenitor-like states and clinical efficacy remains strongest in human samples as a biologically plausible but substantially correlative framework, even though experimental models support a causal role for TCF1/TCF7+ tumor-specific CD8+ reservoirs in sustaining antitumor immunity [44,46,65,75,76,80,81,84].

Physical recovery of T cell–tumor or T cell–APC conjugates is also likely to be most tractable in inflamed, T cell-rich tumors such as melanoma, and more challenging in desmoplastic or immune-excluded epithelial cancers in which T cells are sparse, spatially heterogeneous, or retained in stroma [98–100]. The broader opportunity therefore lies not in assuming that all tumors will yield physical tumor-cell doublets, but in generalizing the engagement principle across antigen-presenting contexts, including autologous APCs, tumor-draining lymphoid compartments, and blood-derived antigen-responsive cells. For these reasons, adhesion should be treated as an organizing and enrichment principle, not as a substitute for antigen identification, native tumor recognition, functional validation, or engineering. The same caution applies to transcriptional signatures. CXCL13, PDCD1, ENTPD1/CD39, TIGIT, tissue-residence programs, and exhaustion-like states can enrich for tumor-reactive cells, and computational tools such as predicTCR and TRTpred can further prioritize candidate tumor-reactive receptors [41,42]. But neither marker expression nor transcriptional prediction is equivalent to tumor specificity unless linked to TCR sequence, antigen recognition, tumor-cell recognition, or clinical function. More importantly for therapy, these approaches usually identify candidates to reconstruct or sort, whereas engagement-based recovery seeks to isolate viable cells that have already converted recognition into productive cellular interaction.

The decisive test of this framework is prospective and comparative. Starting from the same patient material, an engagement-first workflow should recover clonotypes with greater tumor overlap, stronger evidence of neoantigen or tumor recognition, less terminal differentiation, and better post-expansion persistence than expansion-first TIL culture, marker-gated recovery, or peptide-stimulation alone. The decisive endpoint is not enrichment in isolation, but whether selection before expansion produces a cellular product with more relevant clonotypes, better developmental state, and greater capacity for tumor control.

### Therapeutic implication: select first, expand selectively, then reinforce

The therapeutic implication is clear. Engagement-based selection should define the input population, controlled expansion should preserve the selected clonotypes, and antigenic reinforcement should sustain them after transfer. The resulting therapeutic architecture is a matched recovery-and-reinforcement system: patient-specific antigenic information is used not merely to nominate targets, but to recover viable tumor-engaged cells, preserve them through expansion, and sustain them after transfer. Vaccines can induce or expand tumor mutation-specific T-cell responses, but antigen nomination does not by itself establish that the responding clonotypes have productively engaged native tumor in vivo, traffic to the relevant site, resist suppression, or occupy a differentiation state capable of sustaining tumor control in established disease. In other words, the fact that vaccination can expand T cells that recognize a tumor-specific mutation does not necessarily mean that the mutation is an immunologically relevant neoantigen, or that the responding T cells are therapeutically tumor relevant.

Adoptive cell therapy addresses the immediacy problem by delivering large numbers of empirically tumor-engaged T cells rather than waiting for endogenous responses to expand under suppressive tumor conditions. But transferred cells may contract without continued cognate support. Programmable mRNA antigen delivery offers one plausible implementation of this architecture, because the same patient-specific tumor-antigen representation could be deployed across the workflow: as a recovery reagent, a manufacturing guide, and an in vivo reinforcement signal for the transferred product [101,102].

Put differently, the central distinction is between inducing the T cells we wish to have rather than recovering the ones that nature has already provided. Prediction can assist that process, but it cannot substitute for empirical recovery. Productive target adherence is therefore not merely a readout of tumor recognition; it is an enrichment and selection principle that connects specificity, cellular state, manufacturing, and therapeutic amplification. The resulting therapeutic logic is simple: recover the patient’s empirically tumor-engaged repertoire, expand it before irrelevant cells dominate, and reinforce it with the antigenic information that selected it in the first place.

## Methods

### Study design and source selection

This manuscript presents a secondary analysis and synthesis of published studies addressing tumor-reactive T-cell recovery, phenotype, clonotype structure, and functional state. We focused on studies in which tumor-reactive T cells could be evaluated through at least one of the following dimensions: physical interaction with tumor cells or antigen-presenting cells; antigen-dependent recovery, activation, or enrichment; validated neoantigen or tumor recognition; TCR sequence overlap between blood and tumor compartments; or transcriptional evidence of progenitor-like versus terminal cytotoxic differentiation. Studies were included when they enabled direct comparison of tumor-reactive T-cell recovery, clonotype structure, antigen validation, physical engagement, or transcriptional state; they were not selected by formal systematic-review or meta-analysis criteria. The principal analytical anchors were studies of heterotypic T-cell clustering, circulating NeoPBL, NeoSelect-based blood recovery, validated neoantigen-reactive TIL states, and expansion-first TIL or neoantigen-stimulated manufacturing workflows.

### Definitions and analytical framework

For the purposes of this analysis, engagement was defined as stable, antigen-dependent physical interaction between a T cell and a tumor cell or antigen-presenting cell, or recovery of viable T cells through an assay in which antigen-dependent cell-cell interaction was the proximal enrichment step. This included heterotypic T cell–tumor or T cell–APC clustering, stable doublet formation, target-cell extraction after antigen-dependent interaction, and short-term antigen-triggered recovery of viable T cells. Engagement was distinguished from TCR binding alone, activation-marker expression alone, cytokine release alone, proliferation alone, peptide–MHC prediction, or computational prediction of TCR specificity.

### Reanalysis and figure generation

Published figures, tables, supplementary datasets, and statistics were examined to compare tumor-reactive T-cell populations across tumor and blood compartments. When source data were available, enrichment values, differential-expression patterns, clonotype distributions, and gene-expression programs were replotted or reorganized to emphasize common axes of comparison, including physical engagement, clonotype overlap, antigen validation, progenitor-associated programs, checkpoint-associated programs, and cytotoxic-effector differentiation. No new patient samples were collected, and no primary experimental data were generated for this manuscript. Raw sequencing files and single-cell matrices were not harmonized across studies; instead, clonotype distributions, enrichment statistics, transcriptional programs, and functional annotations were compared using engagement and developmental state as common interpretive axes.

### Gene-set and transcriptional-state interpretation

Transcriptional states were interpreted using gene programs commonly associated with progenitor or stem-like CD8+ T-cell states, checkpoint or activation states, and cytotoxic-effector differentiation. Progenitor-associated features included genes such as TCF7, IL7R, KLF2, SELL, CD27, and CD28, whereas cytotoxic-effector features included canonical cytolytic and effector-associated transcripts such as GZMB, PRF1, NKG7, IFNG, and TNF where reported. These programs were used as interpretive axes rather than as rigid cell-state definitions, because tumor-reactive T cells differ across tumor type, anatomical compartment, treatment exposure, and assay format.

### Limitations of secondary analysis

This reanalysis depends on published datasets generated using different experimental platforms, tumor types, patient cohorts, and analytical pipelines. The comparisons are therefore intended to identify convergent biological patterns rather than to provide a pooled quantitative meta-analysis. Physical engagement, antigen-dependent recovery, transcriptional state, and clonotype overlap each provide partial evidence of tumor reactivity, but none alone proves native tumor recognition or clinical efficacy. The conclusions should therefore be tested prospectively in matched workflows comparing engagement-first recovery with expansion-first culture, marker-based sorting, and peptide-stimulation approaches using the same patient material.

## Data availability

All analyses were based on published data cited in the manuscript. No new primary sequencing, clinical, or patient-level datasets were generated. Source values used for figure generation are available in the cited publications and associated supplementary materials where provided. No new human-subject research was performed, and no identifiable patient-level data were analyzed.

## Author contributions

NPR conceived the manuscript, performed the primary analysis and interpretation, and wrote the initial draft. SKV contributed to figure preparation and data visualization. IM and DBT provided scientific comments and manuscript feedback. NDK and DM reviewed the manuscript. All authors approved the final version.

